# Optimisation of a Novel Bio-Substrate as a Treatment for Atrophic Age-Related Macular Degeneration

**DOI:** 10.1101/2019.12.20.884635

**Authors:** Rachel McCormick, Ian Pearce, Stephen Kaye, Atikah Haneef

## Abstract

Atrophic age-related macular degeneration (AMD) is the most common form of AMD accounting for 90% of patients. During atrophic AMD the waste/exchange pathway between the blood supply (choroid) and the retinal pigment epithelium (RPE) is compromised. This results in atrophy and death of the RPE cells and subsequently the photoreceptors leading to central blindness. Although the mechanisms behind AMD are unknown, the growth of fatty deposits known as drusen, have been shown to play a role in the disease. There is currently no treatment or cure for atrophic AMD. Much research focuses on developing a synthetic substrate in order to transplant healthy cells to the native Bruch’s membrane (BM), however, the diseased native BM and related structures still leave the potential for transplanted cells to succumb to disease. In this work we electrospun poly(ethylene terephthalate) (PET) to fabricate a nanofibrous cytocompatible synthetic BM. The apical surface of the membrane was cultured with ARPE-19 cells and the basal surface was decorated with poly(lactic acid-co-glycolic acid) (PLGA) or poly(glycolic acid) (PGA) degradable nanoparticles by electrospraying. The membrane exhibited hydrophilicity, high tensile strength and structurally resembled the native BM. ARPE-19 cells were able to form a monolayer on the surface of the membrane and no cell invasion into the membrane was seen. The presence of both PLGA and PGA nanoparticles increased ARPE-19 cell metabolism but had no effect on cell viability. There was a decrease in pH of ARPE-19 cell culture media 7 days following culturing with the PLGA nanoparticles but this change was eliminated by 2 weeks; PGA nanoparticles had no effect on cell culture media pH. The fluorescent dye FITC was encapsulated into nanoparticles and showed sustained release from PLGA nanoparticles for two weeks and PGA nanoparticles for 1 day. Future work will focus on encapsulating biologically active moieties to target drusen. This would allow this novel bioactive substrate to be a potential treatment for atrophic AMD that would function two-fold: deliver the required monolayer of healthy RPE cells to the macula on a synthetic BM and remove diseased structures within the retina, restoring the waste/exchange pathway and preventing vision loss.

## Introduction

Age-related macular degeneration (AMD) is a progressive disease of the retina that is the leading form of blindness in developed countries. It is a form of central blindness that mainly affects people over the age of 50 years. Although the pathology of AMD is currently unknown, age is considered to be the main contributing factor to the manifestation of this disease (Nowak 2006, Curcio and Johnson 2013). There are two forms of AMD; neovascular and atrophic. Neovascular (or wet) AMD makes up 10% of all reported cases where new abnormal leaking blood vessels break through a layer underlying the retina called the Bruch’s membrane (BM) leading to a loss of central vision. There has been significant developments and improvements in the management of neovascular AMD such as intravitreal injections of anti-vascular endothelial growth factor (anti-VEGF) to prevent any further growth of abnormal blood vessels. Atrophic (or dry) AMD, however, makes up 90% of all cases. It is associated with a slowly progressive form of sight loss where fatty deposits known as drusen, form on BM leading to alterations and atrophy of the RPE. The RPE is the main source of nutrient/waste exchange; if the RPE fails it leads on to photoreceptor death, ultimately leading to central blindness. Although several risk factors for atrophic AMD are known, there are no available treatments other than supportive optical aids.

Much research focuses on developing a synthetic cell transplantation substrate in order to transplant healthy RPE cells onto BM as a treatment for atrophic AMD (Lu, Zhu et al. 2012, Liu, Yu et al. 2014, Surrao, Greferath et al. 2017, da Cruz, Fynes et al. 2018, Tan, Sing et al. 2019). The underlying native diseased BM and related structures remain, however, which leaves the potential for healthy transplanted cells to eventually succumb to disease (White and Olabisi 2017).

Recent research has explored using an anti-inflammatory, antiatherogenic peptide L-4F to reduce the accumulation of fatty deposits on BM of *Macaca fascicularis*, via intravitreal injection of L-4F (Rudolf, Curcio et al. 2019). Those eyes injected with L-4F were found to have had clearance of the fatty deposits along the BM without harming surrounding ultrastructure in the retina. This is particularly interesting, as they described the drug to be well tolerated at even the highest dose, with the only adverse events attributed to the physical process of the injection. Interestingly they found that the effect of the drug was seen bilaterally, even though only one eye was injected with the drug while the other was meant to serve as a control (Rudolf, Curcio et al. 2019).

There is, therefore, scope to provide a two-pronged approach for a potential treatment for atrophic AMD. That is, a persistent bioactive substrate that would function two-fold; provide a permanent basement layer for transplantation of healthy RPE cells to the area required, while removing diseased structures (drusen) on the BM and fatty deposits (lipofuscin) within the retina (Curcio and Johnson 2012), using moieties such as L-4F. This would replenish the native BM nutrient/waste exchange pathway, preventing the progressive loss of photoreceptors, through a more controlled and localized effect. Our aim is to develop a bioactive persistent cell transplant substrate for the treatment for atrophic AMD. This article describes the development of a composite membrane for this two-pronged approach.

## Material and methods

### Electrospinning of PET scaffold

Poly(ethylene terephthalate) PET pellets (04301, Polysciences Inc) were dissolved in neat 1,1,1-3,3,3-hexafluoroisopropanol (HFIP) (Apollo Scientific Ltd.) at a concentration of 17.5% (w/v). The fibres were collected for 15 minutes under laboratory conditions on a grounded plate covered with aluminium foil at a working distance of 15cm, flow rate of 2ml/h and a voltage of 25kV.

### Electrospraying of PLGA and PGA nanoparticles

Poly(lactic acid-co-glycolic acid) 50/50 (PLGA) (26269, Polysciences Inc) was dissolved in neat chloroform (Merck) at a concentration of 2% (w/v). Poly(glycolic acid) (PGA) (06525, Polysciences Inc) was dissolved in neat HFIP at a concentration of 1% (w/v) and left to stir overnight at room temperature. For nanoparticle encapsulation, 2mM of FITC was added to the solvents at the same time as the polymers. The homogeneous polymer solution was introduced into a 10mL plastic syringe and a blunt-tip needle attached. Any air bubbles were removed and the filled syringe was fixed in a mechanical syringe pump. All work was carried out at ambient conditions in a fume-hood. Nanoparticles were sprayed into a glass dish filled with 0.1% (v/v) isopropanol (Merck) in dH_2_O placed on a magnetic stirrer, connected to a grounding plate and left to spray for 1h. Working parameters for PGA nanoparticles were 25kV, 25cm working distance, 2ml/hr flow rate; and for PLGA nanoparticles; 11kV, 13cm working distance, 0.5ml flow rate.

### Fibre and nanoparticle diameter measurements

The diameter of 50 fibres and nanoparticles from 3 different images of the electrospun fibres and electrosprayed nanoparticles were measured using ImageJ. (n = 3).

### Preparation of fibres and nanoparticles for SEM

Electrospun membrane or 10μl of electrosprayed nanoparticles suspended in solution was placed on a carbon tab (TAAB) mounted on an aluminium stub (TAAB). The nanoparticles were surrounded by a layer of silver dag (Merck) and were left overnight in a desiccator for the solution to evaporate. Membrane or nanoparticles were gold sputter coated (Quorum) and imaged using SEM (Quanta FEG250 ESEM) with EHT of 5kV. (n = 3).

### Contact angle measurements

Electrospun membrane was cut into 3cm × 1cm rectangles. Samples were either untreated, UV treated (1 hour), placed in ethanol, or placed in cell culture medium and left to dry before being measured for changes in wettability using DSA 100 (Kruss-Scientific). (n = 6).

### Tensile testing

Quantitative tensile testing of the electrospun membrane was undertaken using UniVert tensile tester (CellScale) equipped with a 10N load cell at a displacement rate of 12mm/minute. The membranes were cut into dog-bone shaped strips 2cm in length by 0.5cm in width and tested until failure or until the tensile tester had reached maximum distance (n = 16). Membrane thickness was measured using a digital micrometer (HITEC, 190-00, Farnell). For wet samples (n = 7), samples were soaked in dH_2_O before mounting into the tensile tester.

### FTIR

Electrospun membranes were cut into 3cm × 1cm rectangles. Samples were either untreated, UV treated (1 hour), placed in ethanol, or placed in cell culture medium and left to dry before being measured using Vertex 70 Fourier Transform Infrared Spectrometer (Vertex). (n = 6).

### Cell culture

Electrospun membrane was cut into 1.5cm^2^ squares and placed in Scaffdex (Merck). ARPE-19 cells (ATCC-LGC, CRL-2302) were seeded at a density of 50,000 cells/sample and incubated at 37°C, 5% CO_2_, 98–99% humidity and grown for 1 month or 3 months. Media consisted of DMEM:F12 (Merck) containing 10% FCS (Thermofisher Scientific), 2.5mg/L amphotericin B (Merck) and penicillin-streptomycin (Merck). Media was changed every three days. Controls were glass coverslips (Agar).

For long-term culture, the membrane was fabricated as aforementioned and then mounted onto a grounded water bath as described. Nanoparticles were sprayed onto the membrane using the described parameters followed by cutting and mounting into Scaffdex with the nanoparticle decorated side orientated to be the basal side. Cells were seeded onto the apical side and cultured as mentioned above.

### Histology

Cell cultured membranes were fixed in 10% NBF for 15 minutes and processed for histological staining with Leica TP1020. Following processing, tissues were embedded for sectioning in paraffin wax and stained with H&E using standard protocol. (n = 4).

### Preparation of cells for SEM

Cells were fixed in 1.5% Glutaraldehyde (Fluka) for 30 minutes at 4°C and dehydrated with graded ethanol; 2 × 3 min 50%, 2 × 3 min 70%, 2 × 3 min at 90%, 2 × 5 min at 100%. Hexamethyldisilane (HMDS, Merck) was added for 2 × 5 mins and left overnight to evaporate. Samples were mounted on a carbon tab on an aluminium stub, gold sputter coated (Quorum) and imaged using SEM (Quanta FEG250 ESEM) with EHT of 5kV. (n = 3).

### Immunofluorescence staining

Cell cultured membranes were harvested after 1 or 3 months of culture, washed with DPBS and fixed in 10% NBF for 15 minutes. Cells were permeabilised with 1% Triton-X for 10 minutes, blocked with 10% goat serum for 30 minutes and incubated with primary antibodies (table 1) overnight at 4°C. The following day, cells were incubated with the appropriate secondary antibody for 1 hour, followed by DAPI staining, and membrane placed on glass slides and mounted to coverslips with Vectorshield (Vectorlabs). Slides were viewed using the Zeiss ImagerM1 microscope. (n = 3).

**Table 1.** Primary antibody stains and the companies from where they were bought with corresponding catalogue numbers.

| Primary antibody | Company (catalogue number) |
| --- | --- |
| Anti-Bestrophin/BEST1 antibody | Abcam (ab14927) |
| Anti-RPE65 antibody | Abcam (ab13826) |
| Anti-LRAT antibody | Abcam (ab166784) |
| Anti-PEDF antibody | Abcam (ab180711) |
| Anti-CRABP1 antibody | Abcam (ab235838) |
| Anti-MERTK antibody | Abcam (ab110108) |
| Anti-LAMP2 antibody [H4B4] | Abcam (ab25631) |
| Anti-USO1 antibody | Abcam (ab102470) |
| Anti-GULP antibody | Abcam (ab236893) |
| ZO-1 | Thermofisher Scientific (61-7300) |
| Alexa Fluor 488 Phalloidin | Thermofisher Scientific (A12379) |

### Resazurin Assay

Resazurin solution (5mg resazurin salt in 40mL Dulbecco’s phosphate buffered saline) (DPBS) (PAA laboratories), was added to each sample (100µl/ml) and returned to the incubator for 4 hours. A 100μl aliquot of media was taken from each sample and transferred into a black bottomed 96-well plate. Fluorescence was measured at 530–510nm excitation and 590nm emission using a fluorescence reader (FLUOstar OPTIMA, BMG LABTECH). (n = 4).

### Barrier assay

Scaffdex mounted membranes, with or without cells, were placed in 12 well plates with DPBS on the basal side of the membranes. On the apical side of the membranes, 1mM FITC (Merck) in DPBS was added. Every 10 minutes, 100µl of DPBS was taken from the basal side of the membranes and measured using the fluorescence reader (FLUOstar OPTIMA, BMG LABTECH). (n = 4).

### Live/dead assay

Cell cultured membranes were washed in DPBS and following manufacturer’s instructions (molecular probes) incubated with calcein AM and ethidium homodimer-1 for 30 minutes. The material was placed on glass slides and mounted to coverslips with Vectorshield (Vectorlabs). Slides were viewed using the Zeiss ImagerM1 microscope. (n = 3).

### pH measurements

Three pH measurements were taken from each well using a digital pH meter (Mettler-Toledo). (n = 6).

### Measurement of nanoparticle degradation

Following electrospraying, 10ml of collected nanoparticles in 0.1% (v/v) isopropanol was kept at 37°C. An aliquot (1ml) of solution was then collected at each time point and kept at −20°C until the final time point. For collection from cell cultures 1ml of media was collected at each time point and media changed. A 100μl aliquot was taken from each sample and transferred into a black bottomed 96-well plate. Fluorescence was measured at 530–510nm excitation and 590nm emission using a fluorescence reader (FLUOstar OPTIMA, BMG LABTECH). (n = 4).

### Statistical analysis

Data are presented as mean (+/− standard error) with n values noted within the text. Analysis was undertaken using GraphPad Prism software using t-tests or one-way ANOVA followed by the appropriate post-test noted within the text. Significance was p<0.05.

## Results & Discussion

### Characterisation of electrospun PET: Morphology, FTIR, hydrophilicity, mechanical properties, barrier assay, and cell culture

Scaffolds were characterized for adequate mechanical properties to undergo surgical handling and the ultrastructure was characterized to match fibre morphology to mimic the native BM (Yamamoto and Yamashita 1989, Del Priore, Tezel et al. 2006). Surface chemistry was analyzed for presence of functional groups following surface treatment and hydrophilic properties measured using water contact angle (WCA). Barrier properties were analyzed with and without a monolayer of cells to ascertain any changes with cell culture.

Nanofibrous non-degradable PET fibres were produced in a collectable membrane form. SEM micrographs showed the membranes exhibited a randomly orientated fibrous mesh (figure 1A); an open network of interconnected voids with average fibre diameter of 387nm (+/−147.8) (figure 1B). The membrane was able to undergo folding, rolling, and twisting without being destroyed; depicting the ease with which it could be handled and exhibiting adequate mechanical properties (figure 1C-D). The handling properties were in agreement with previously published work, even though the concentration of the polymer solution used was lower than previous work (Haneef and Downes 2015). Morphologically this electrospun membrane mimics native BM (Yamamoto and Yamashita 1989).

**Figure 1.**
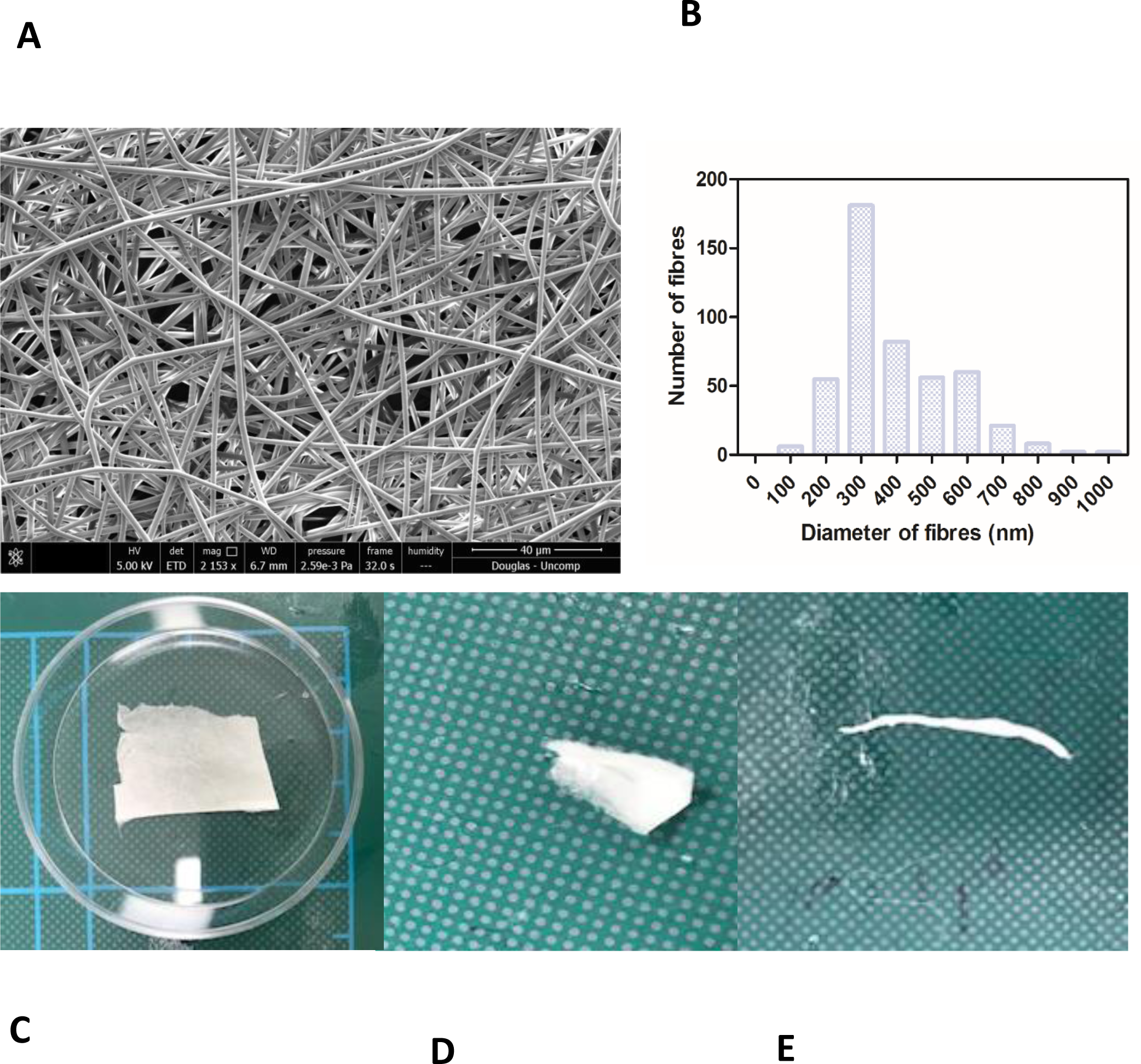
SEM micrograph of 17.5% PET electrospun membrane exhibiting ultrastructure of randomly orientated fibres (**A**) and corresponding histogram showing distribution of fibre diameter size; average diameter of 387nm (+/−6.9) (**B**). Photographs of material folded (**C**), rolled (**D**), twisted (**E**) without breaking denote good handleability of the material (n = 3). Scale shown = 5cm.

The membrane gave an interesting stress vs. strain profile following tensile testing; exhibiting an elastic region (original form is retained following release of load) and a plastic region (membrane deforms indefinitely) before attaining failure exhibiting the ultimate tensile strength (UTS) (figure 2A-B). 16% of membranes tested did not break (figure S1, supplementary data). Soaking the membranes in dH_2_O did not significantly change the Young’s modulus (YM) of the elastic region (figure 2C), however, the YM of the plastic region significantly increased from 5.5MPa (+/−0.8) to 13.8MPa (+/−3.5) and the UTS significantly increased from an average of 11.23MPa (+/−1.5) to 20.29MPa (+/−5.3) (figure 2D-E). Wetting the membrane introduced hydrogen bonding between the fibres thereby increasing the YM of the plastic region and the UTS, while in the dry form the fibres did not have the water to provide the added interactions (Chen, Cheng et al. 2018, Kurokawa, Kimura et al. 2018).

**Figure 2.**
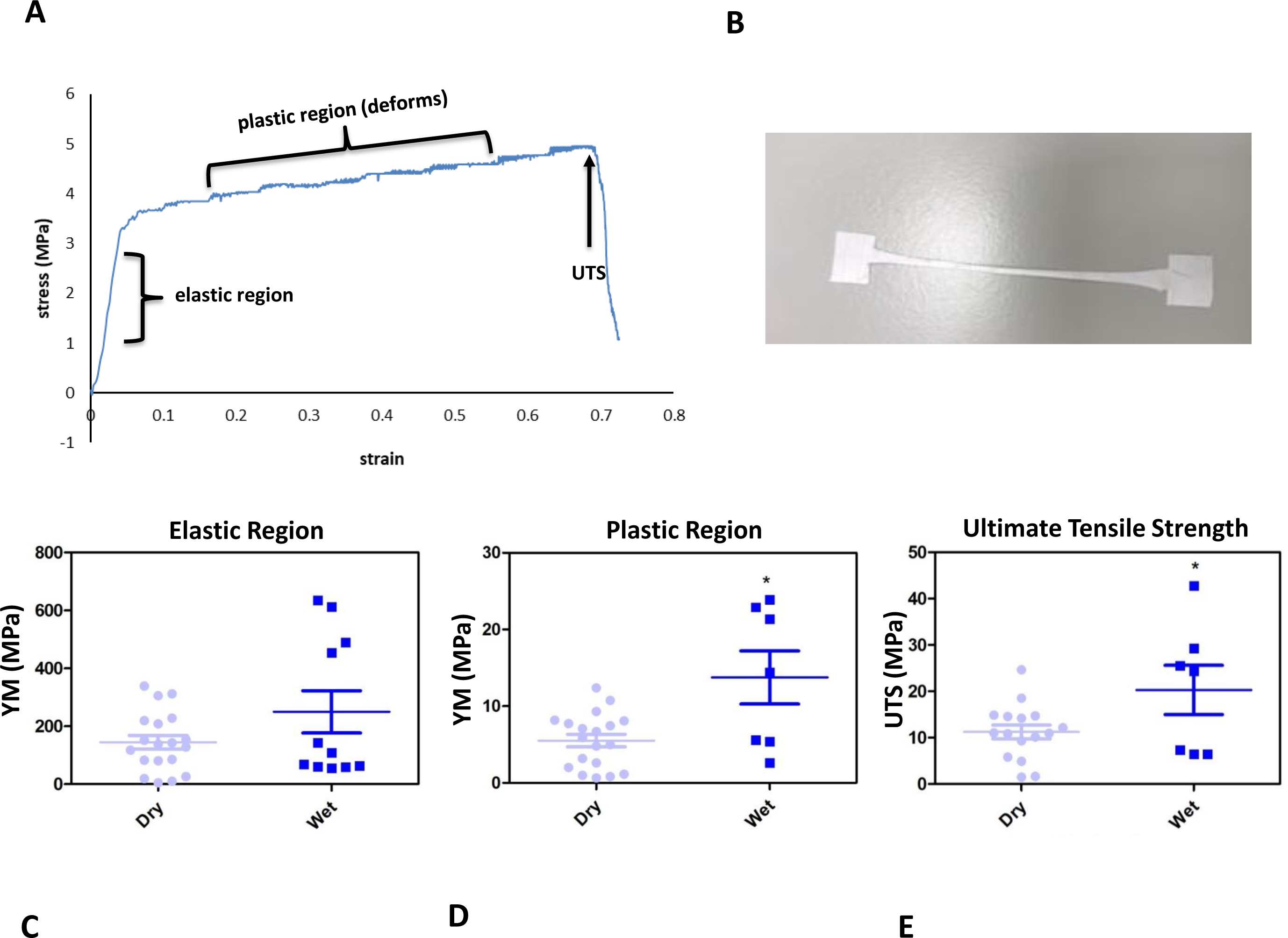
The membranes gave an interesting stress vs. strain plot, as shown by the representative profile (**A**) and stretched to deformation (**B**). Young’s modulus (YM) for the elastic region (retains form) are high, with no significant difference between the dry (145.2 MPa (+/−23.46)) and wet (250.2 MPa (+/−73.08) membranes (**C**), whereas the plastic region (deformation) YM significantly increased from 5.15 MPa (+/−0.82) (dry) to 13.75 MPa (+/−3.47) (wet) (**D**), and the ultimate tensile strength (UTS) of the membrane significantly increased from 11.23 MPa (+/−1.5) (dry) to 20.29 MPa (+/−5.3) (wet) (E). Significant difference *= p<0.05 compared with dry material, t-test (n = 16 for dry, n = 7 for wet). Data presented: mean (+/− standard error).

Treatment of the membrane with cell culture media significantly increased hydrophilicity compared to all other treatment methods, whereas ethanol treatment resulted in an increase in the hydrophilicity compared only to UV treatment. WCA of control membranes averaged at 132.6° which decreased to 85.8° and 118.4° following treatment with media and ethanol respectively (Figure 3A-B, table S1, supplementary data). FTIR spectra exhibit the characteristic carbonyl peak in PET membrane typical of the ester bond group within the polymer structure, denoted by the strong peak at ~1700cm^−1^ for C=O. No changes were detected for any of the membranes except the cell culture media treated membrane. A peak at ~3400cm^−1^ denoting the presence of O-H group was detected, suggesting that the surface of the membrane underwent a degree of functional group opening to form-COOH free groups that could be attributed to protein interaction or hydrolysis of the surface of the polymer (Figure 4A-D) (Haneef and Downes 2015, Kawai, Kawabata et al. 2019).

**Figure 3.**
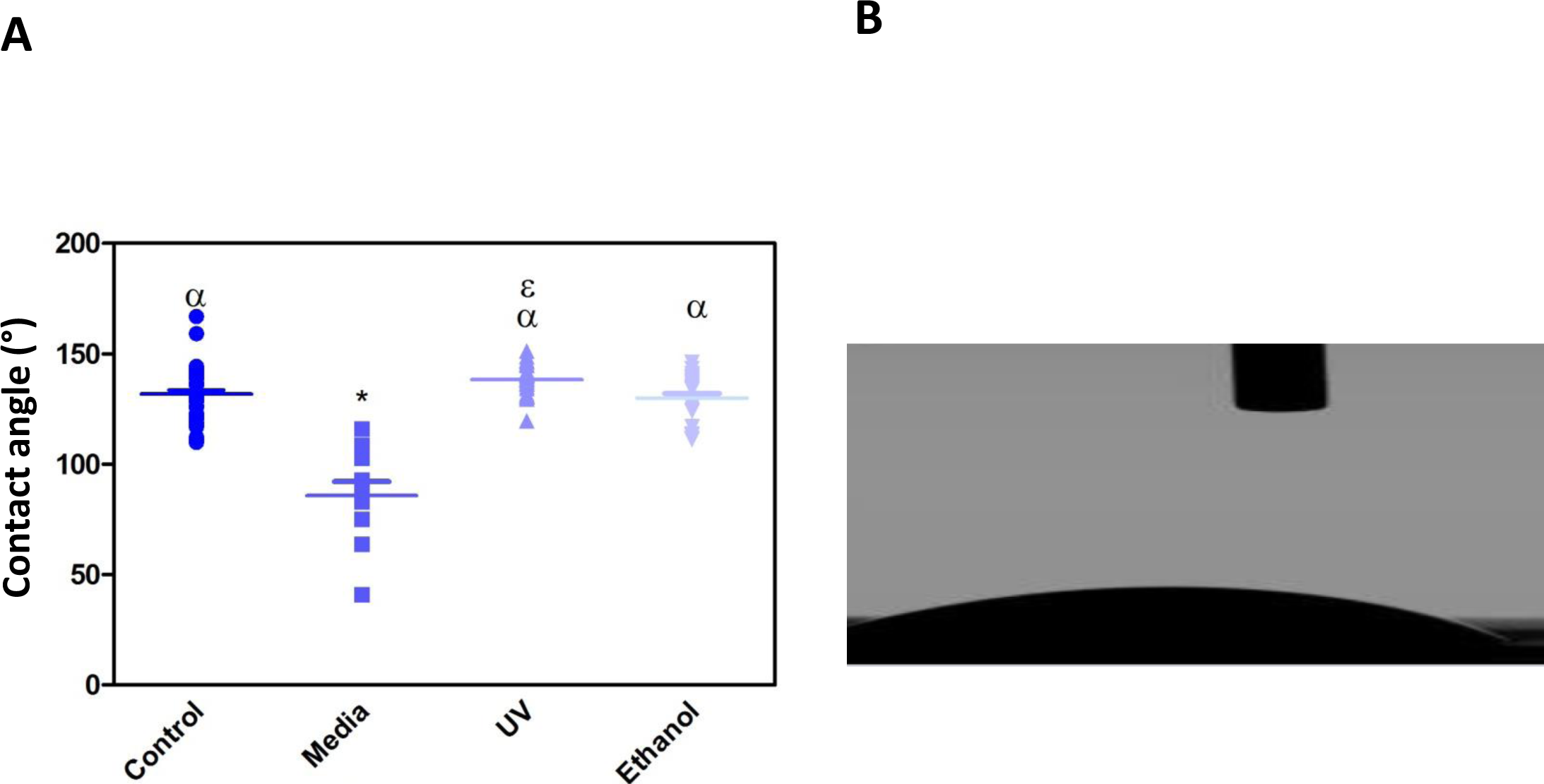
Dot-plot exhibiting the WCA of PET untreated control, cell culture media treated, UV treated and ethanol treated membranes. Media treated membrane exhibited a significantly different decrease in WCA (85.8° +/−20)) compared to control and all other treatments (**A**). One-way ANOVA followed by Tukey’s post hoc test (n = 6). * = p<0.05 significantly different to untreated control, α = p<0.05 significantly different to cell culture media treated, ε = p<0.05 significantly different to ethanol treated. Data presented: mean (+/− standard error). Photograph of WCA show the hydrophilic nature of the material (**B**).

**Figure 4.**
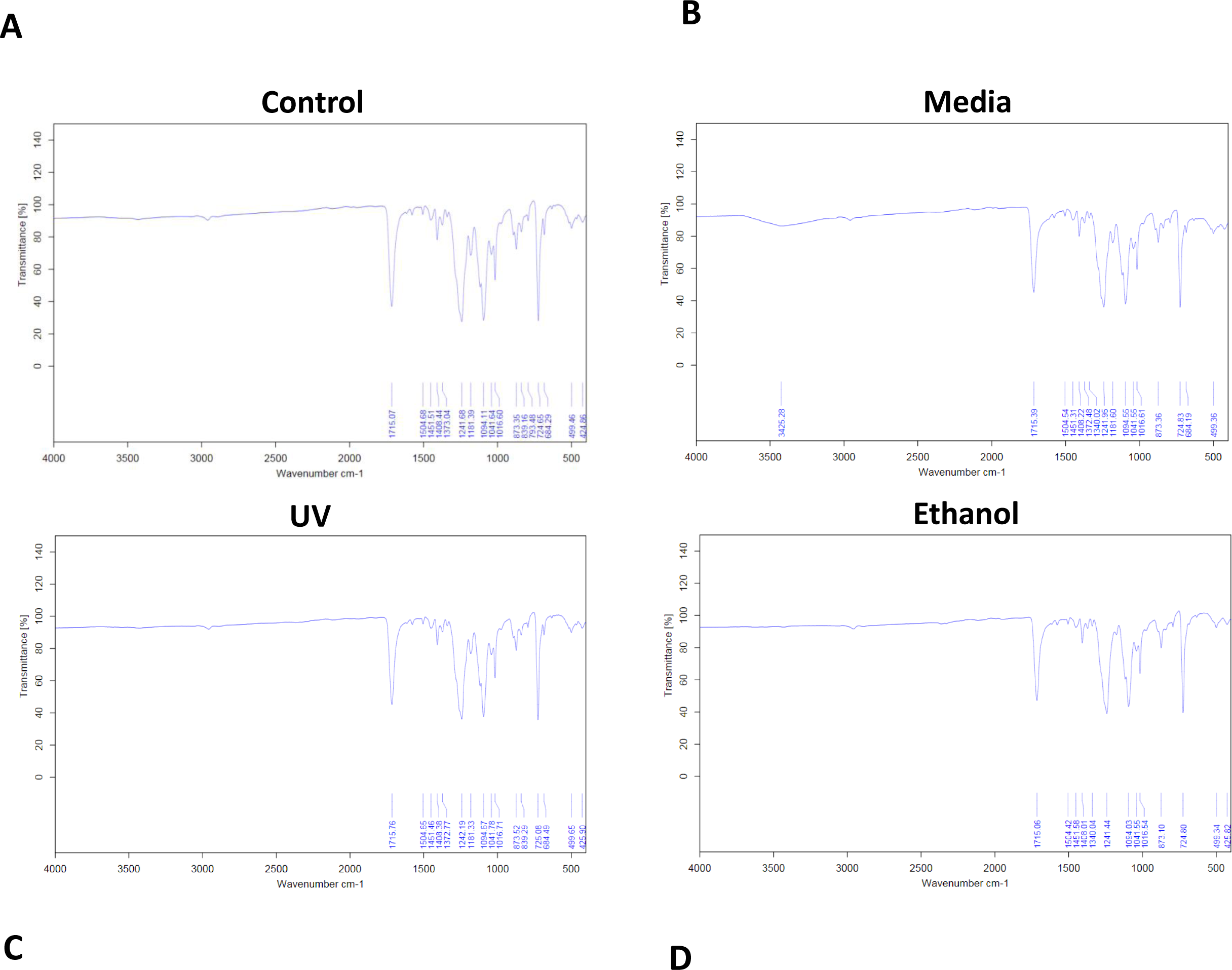
FTIR spectra of electrospun membrane untreated control (**A**), tissue culture media treated (**B**), ethanol treated (**C**) and UV treated (**D**). All spectra exhibit the characteristic PET carbonyl peak at ~1700cm^−1^ with the media treated membrane exhibiting a peak at ~3400cm^−1^ suggesting evidence of protein interaction (n = 6).

SEM and phalloidin staining showed cells were able to form a monolayer when cultured on the membrane up to 3 months, with microvilli that are phenotypical of RPE cells (Thomson, Treharne et al. 2011) presenting after 1-month’s culture (Figure 5A-F). Although the auto-fluorescence of the membrane fibres made it challenging to see clearly, cells plated on the membrane stained positively for the characteristic proteins of RPE cells (Liao, Yu et al. 2010, Brandl, Zimmermann et al. 2014) in comparison with positive controls (figure 6). This is particularly noticeable with BEST1 (bestrophin-1), a Ca^2+^-activated Cl^−^ channel (CaCC) specifically expressed in the retinal pigment epithelium (RPE) of the eye (Kittredge, Ji et al. 2018), USO1 (General vesicular transport factor p115) and GULP (PTB domain-containing engulfment adaptor protein 1), an adaptor protein involved in phagocytosis (Park, Kang et al. 2008), an important function of RPE with involvement in the phagocytosis of photoreceptor outer segments (Sparrow, Hicks et al. 2010).

**Figure 5.**
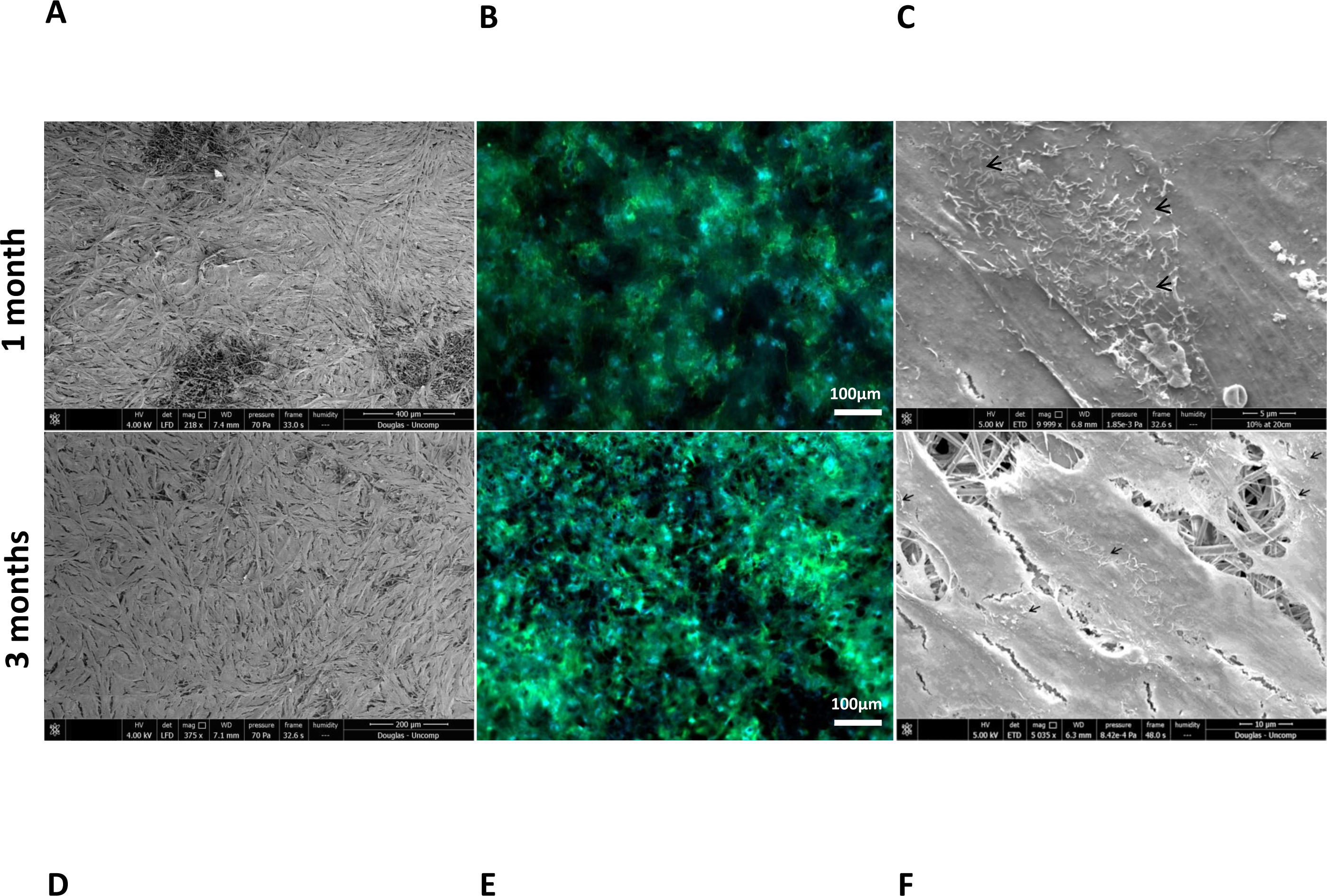
Representative SEM micrographs and corresponding fluorescence images of DAPI/phalloidin stained ARPE-19 cells cultured on PET membranes for 1 month (**A, B, C**) and 3 months (**D, E, F**). Cells populate the membrane with better coverage and more apparent phalloidin staining at 3 months culture. Microvilli (arrows) phenotypical of RPE cells are apparent by 1 month. (n = 3).

**Figure 6.**
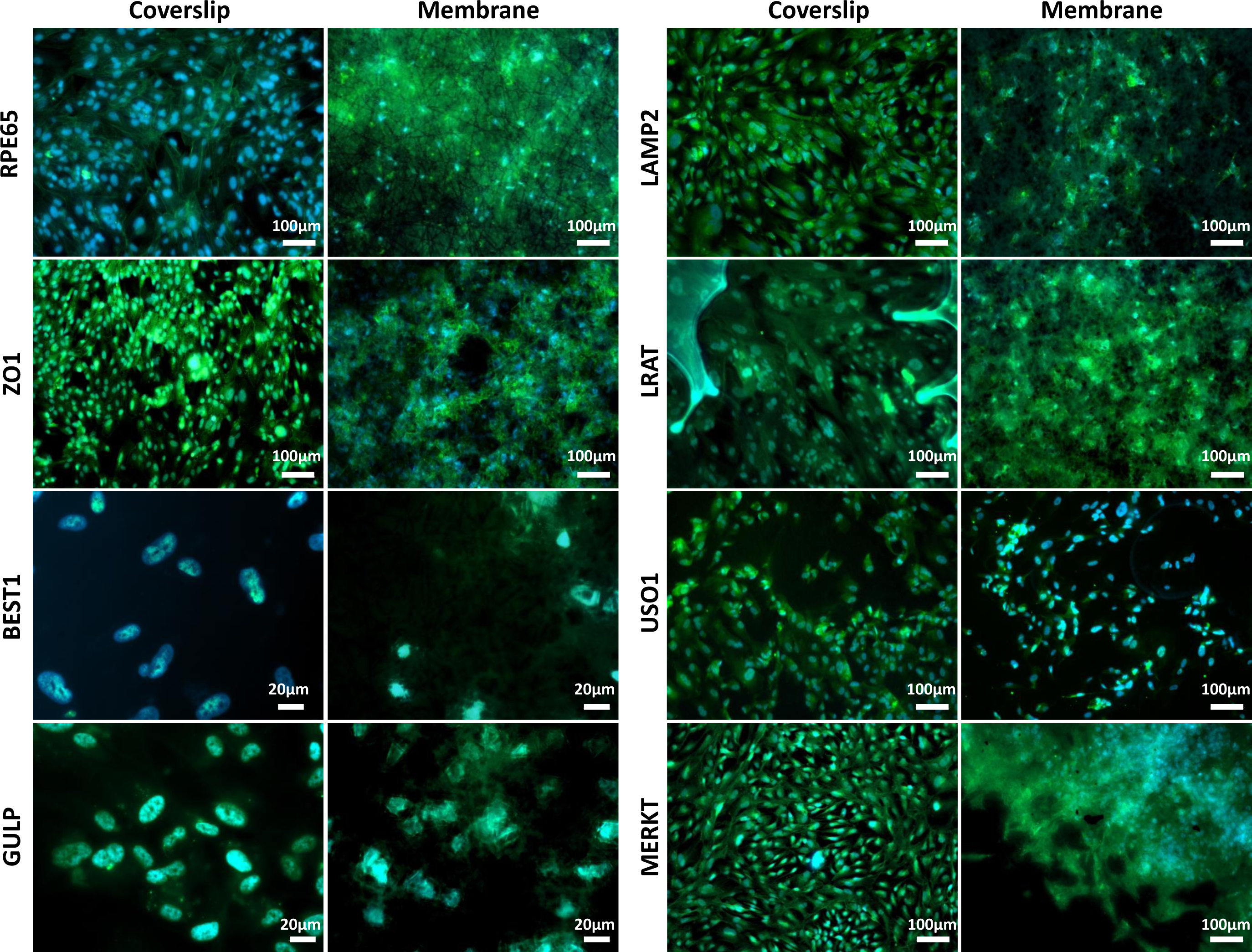
Representative immunofluorescent images show RPE cells cultured on the electrospun membrane for 1 month stained positively for RPE cell marker proteins and RPE function proteins (green). Cells cultured on the membranes were difficult to image clearly due to the auto-fluorescent nature of the PET fibres. Upon comparison with the positive glass coverslip control, similar staining patterns are apparent, especially noticeable with BEST1, GULP and USO1. DAPI = blue nuclear stain (n = 3).

H&E stained sections showed the lack of cell invasion in the bulk structure of the membrane, which was the desired characteristic at both 1 week and 3 months’ culture (Figure 7A-B) This was further confirmed with SEM imaging showing the absence of any cells on the basal side of the membrane following cell culture on the apical side (figure 7C).

**Figure 7.**
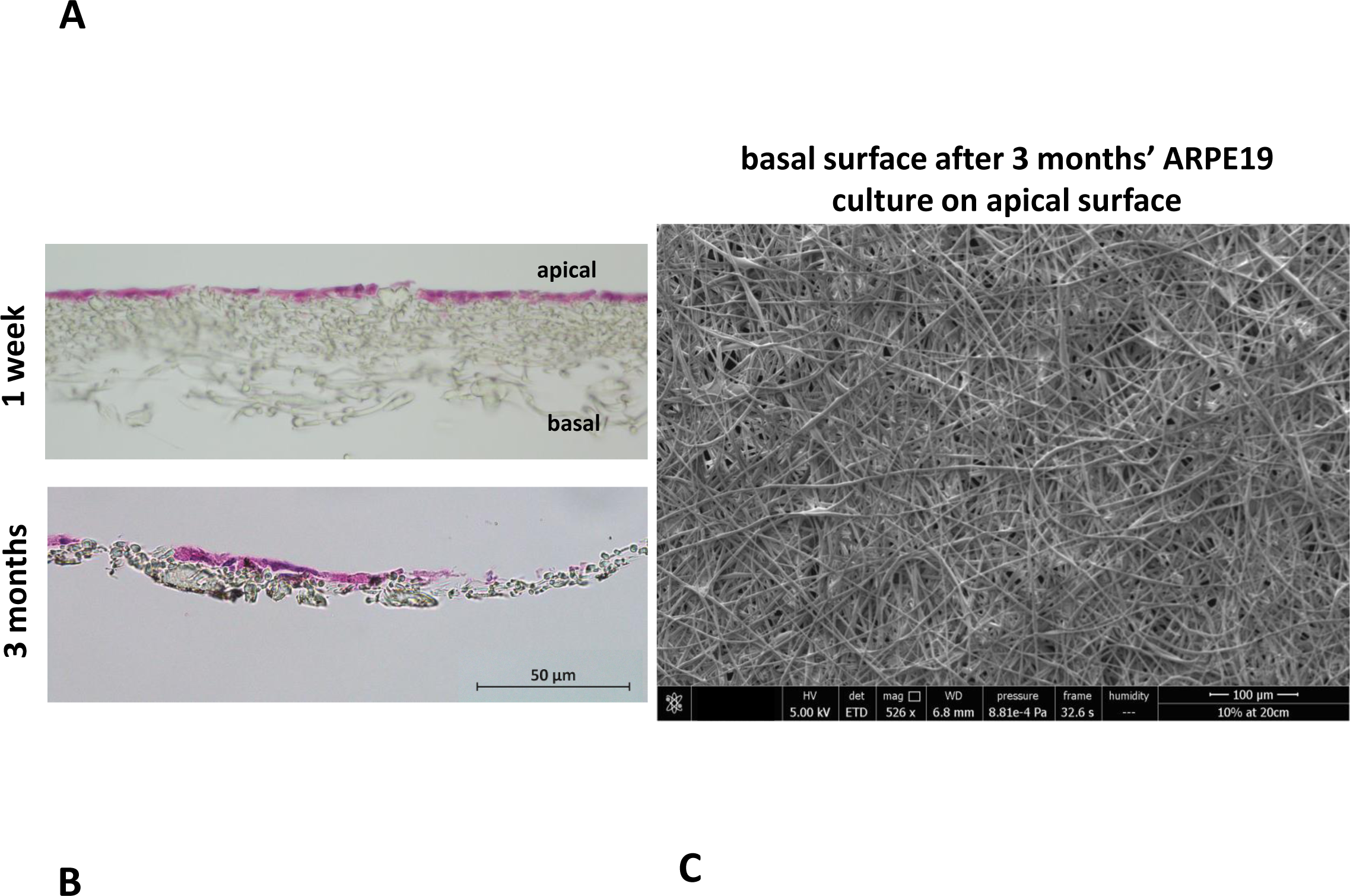
The membrane does not allow invasion of cells through its architecture. H&E staining of cells cultured on the membrane for 1 week (**A**) up to 3 months (**B**) show the desired lack of cell invasion. The SEM micrograph (**C**) exhibits a lack of cells on the basal side of the membrane that had been seeded with ARPE-19 cells on its apical side and cultured for 3 months (n = 4).

Resazurin assays showed the metabolism of the cells cultured on the membrane was significantly lower compared to positive control in the initial first week of culture, thereafter, the metabolism was not significantly different to the positive control (figure 8). This may have been due to cells not adhering as quickly due to the porosity of the membrane, as this difference in metabolism was overcome by 2 weeks. These data are in contrast to previous studies by our group that have shown RPE cells had increased metabolism when plated on an electrospun PET membrane (Haneef and Downes 2015), however, the membrane formed in the previous study was electrospun for 1 hour and had thinner fibres, giving the membrane a different morphology. Current results show that ARPE-19 cells were able to form a monolayer, express proteins and exhibit microvilli phenotypical to the RPE cells, after long-term culture on the current electrospun membrane.

**Figure 8.**
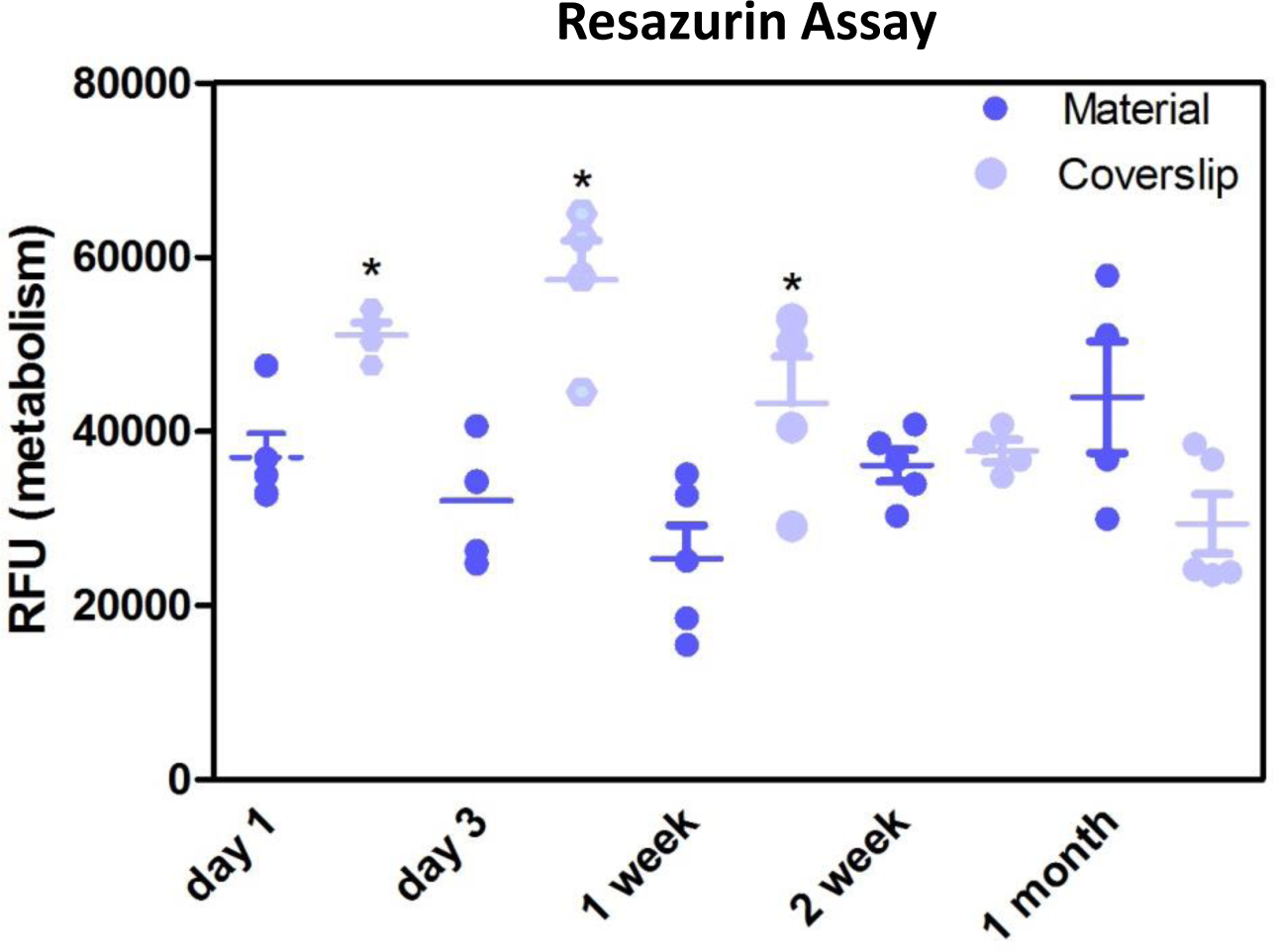
Resazurin assay showed metabolism of cells cultured on the membranes was significantly lower in the initial 1 week of culture compared to glass coverslip controls. There was no significant difference in the metabolism of the cells thereafter up to 1 month. Significant difference *p<0.05, one-way ANOVA followed by Bonferroni post-test (n = 4). Data presented: mean (+/− standard error).

To determine the permeability of the membrane a barrier assay was carried out on acellular membranes and on membranes cultured with ARPE-19 cells for 1 month, as this was when a monolayer had formed. The barrier assay on acellular membranes showed FITC was able to passively diffuse through the membrane in a time dependent manner, taking 80 minutes for maximum signal to be detected (figure 9A). Cell cultured membranes showed the transport of FITC had increased with fluorescence plateauing within the first 10 minutes of culture, suggesting the monolayer of cells were actively transporting the FITC across the membrane and functioning correctly (figure 9B). These data showed that cells plated on the material did not affect the permeability of the membrane and shows it has the appropriate properties to act as an artificial BM.

**Figure 9.**
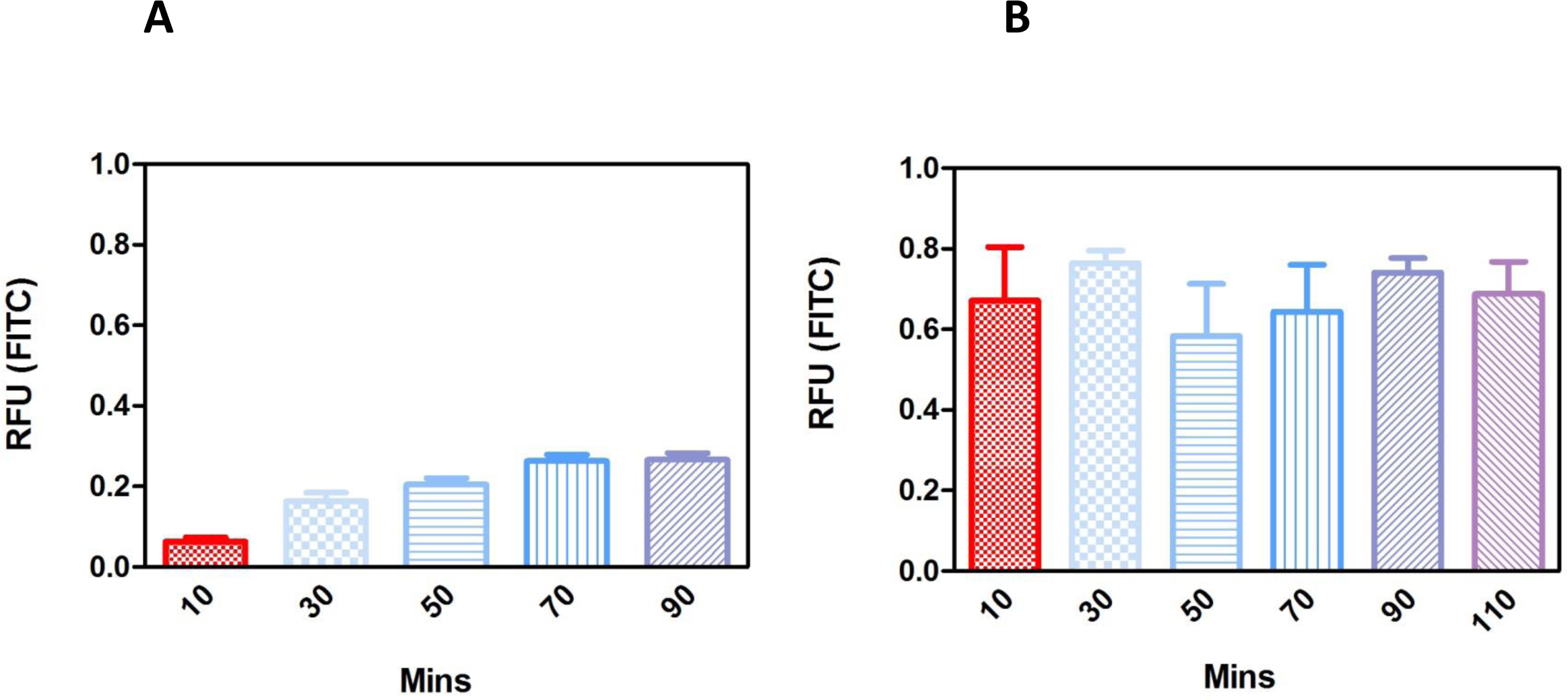
Fluorescence barrier assays in the acellular substrate showed passive diffusion that plateaued FITC concentration across the membrane by 70 minutes (**A**), whereas fluorescence in the 1-month cell cultured membrane plateaued in the first 10 minutes of measurement suggesting the active transport of FITC by the monolayer of cells (**B**) (n = 4). Data presented: mean (+/− standard error).

Future experiments to confirm the ability of the RPE cells to perform phagocytosis when seeded on the PET membrane are needed. The presence of the phagocytosis marker MERKT in the ARPE-19 cells cultured on the PET membrane (figure 6) suggested the cells had phagocytic potential. Furthermore, a study by Cruz et al, have shown the ability of human embryonic RPE cells plated onto a PET membrane to phagocytose both *in vitro* and following surgical implantation (da Cruz, Fynes et al. 2018).

### Nanoparticles

Spherical nanoparticles were successfully fabricated with average nanoparticle diameters at 480nm (+/−252) for PLGA and 59nm (+/−49) for PGA (Figure 10A-D). Interestingly less variation in particle size was seen in PGA particles.

**Figure 10.**
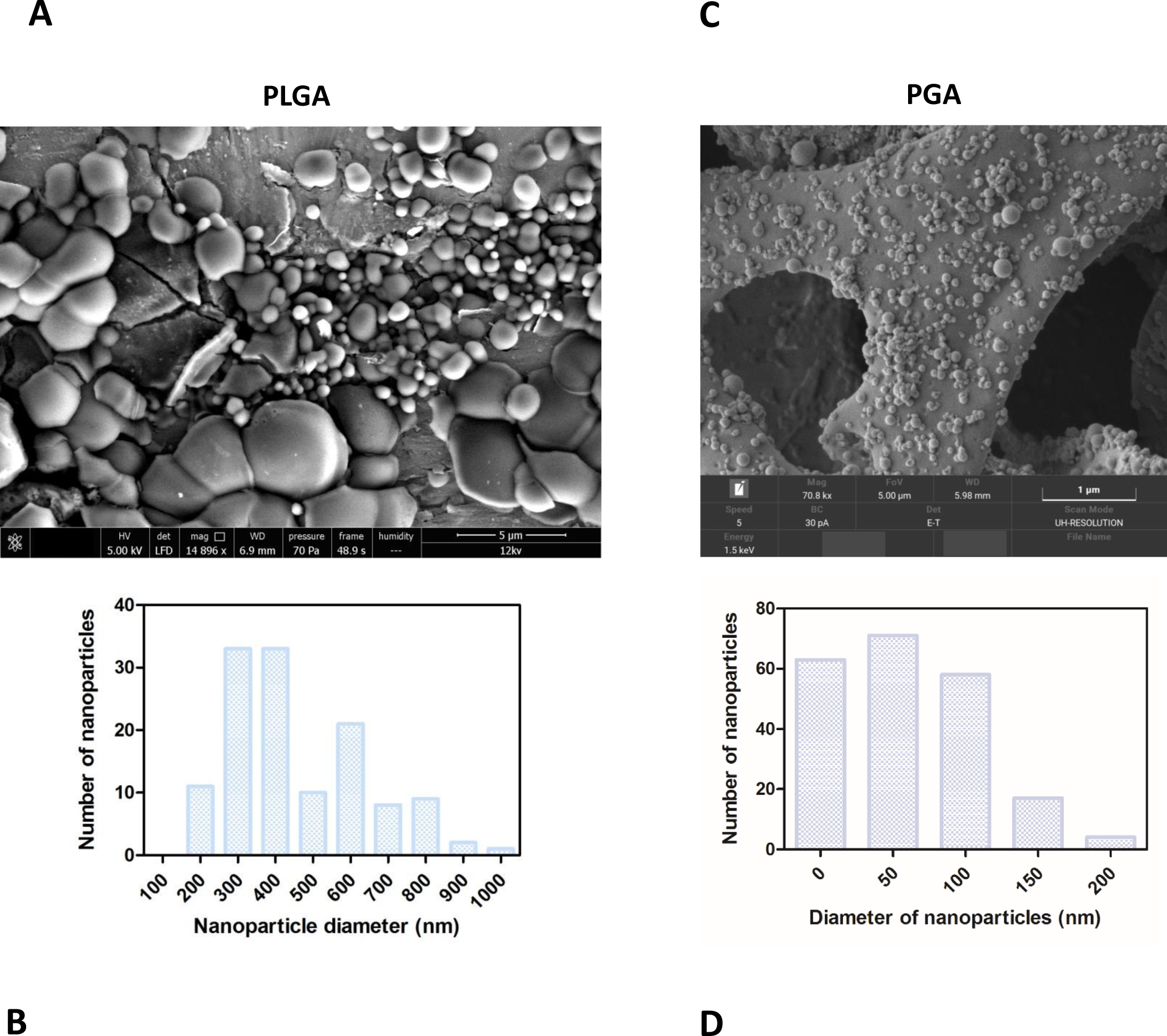
SEM micrographs show PLGA nanoparticles (**A, B**) were larger than PGA nanoparticles (**C, D**) Corresponding histograms showed PLGA averaged at 480nm (+/−252) with a broader range of particle sizes, whereas PGA averaged at 59nm (+/−49) and exhibited less variation in particle size (n = 3). Data presented: mean (+/− standard error).

The nanoparticles did not affect ARPE-19 cell viability or morphology when compared with the membrane only control (figure 11); cells attached 24hours following seeding and formed a monolayer by 1 month in the presence of nanoparticles. Resazurin assays showed the presence of nanoparticles induced a significant increase in cell metabolism compared to the membrane only control up to 2 weeks, which resolved thereafter up to 1 in month in culture (figure 12). The increase in metabolism may be attributed to the change in pH as the nanoparticles degrade with time; it was found that the pH of the culture media decreased after 1 week for both PLGA and PGA nanoparticles (figure 13, table S2, supplementary data). PLGA nanoparticles significantly decreased the pH up to 1 week’s degradation compared to membrane only control. The metabolites of PLGA degradation are lactic acid and glycolic acid, which contribute to lowering the pH of the surrounding environment. It could also be attributed to the larger variation in particle size which could be contributing to the significant change in pH compared to PGA.

**Figure 11.**
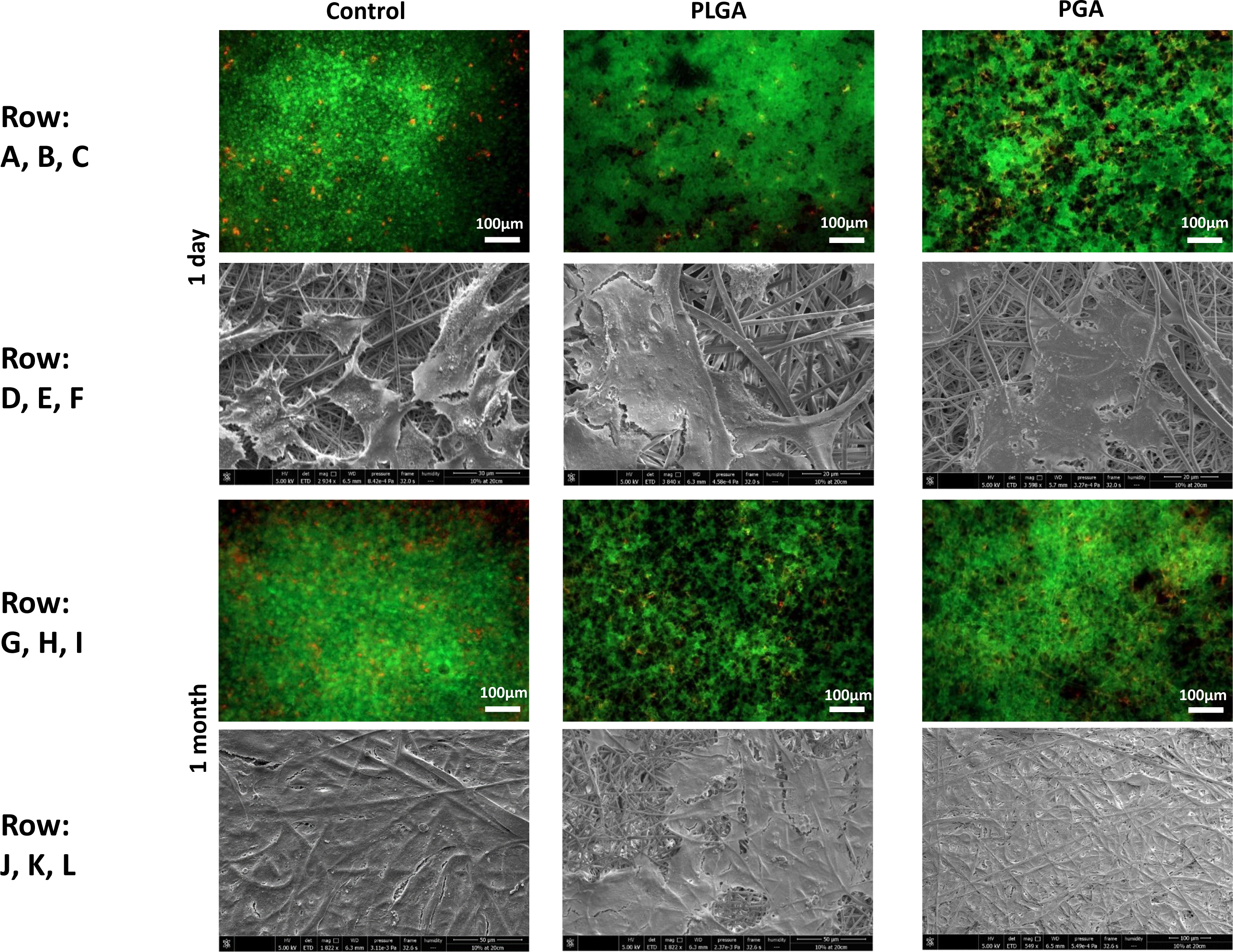
Representative images of live/dead staining (green/red) and corresponding SEM micrographs of ARPE-19 cells after 1 day’s culture (**A-F**), 1 month’s culture (G-L) on membrane only control, or with either PLGA or PGA nanoparticles decorated on the basal side of the membrane. Live/dead images show there is little difference between the three conditions with few dead cells present up to 1 month’s culture. SEM micrographs show the progression in cells covering the membrane’s apical surface (n = 3).

**Figure 12.**
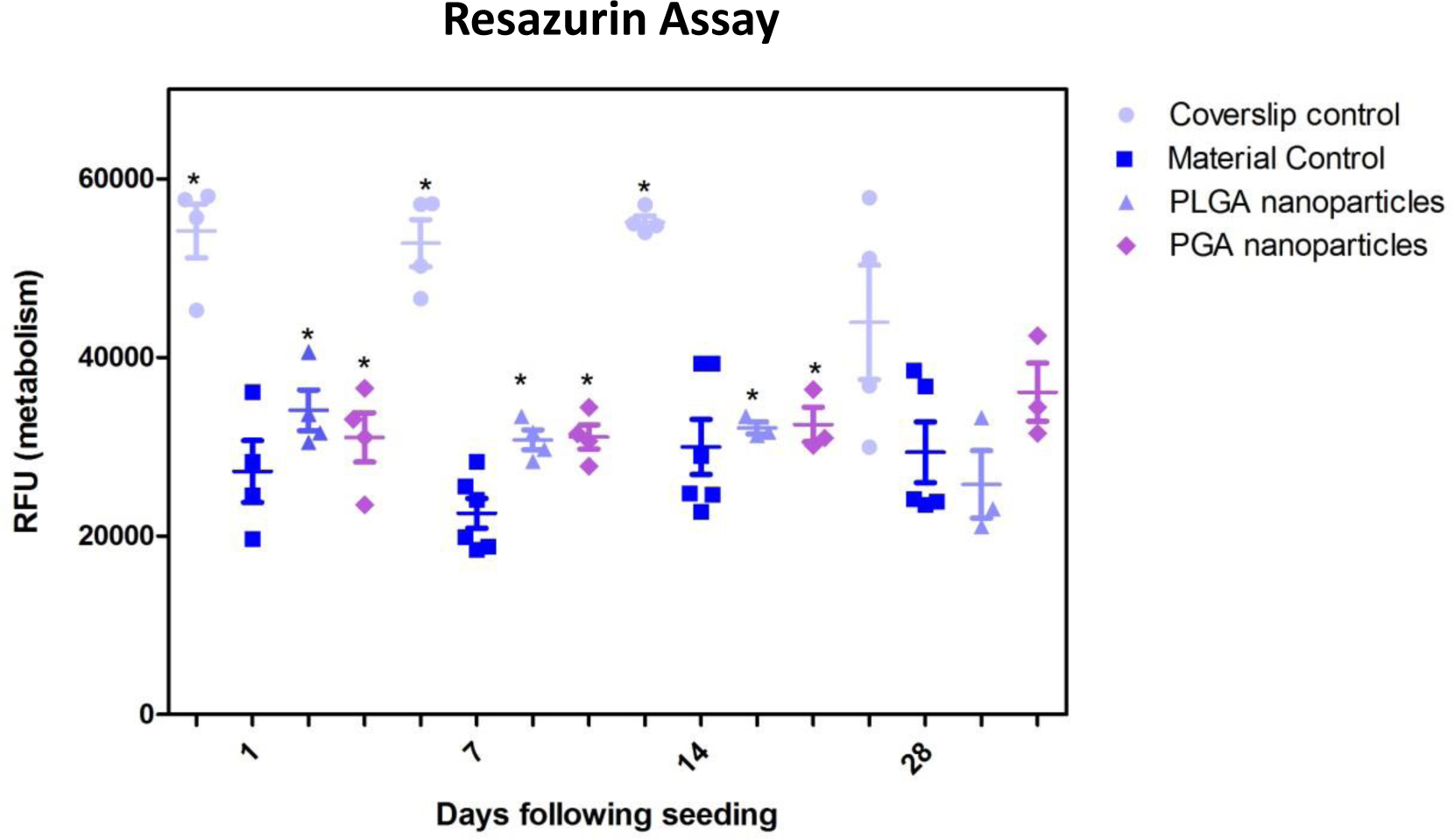
Resazurin assay of ARPE-19 cells plated on membranes with PLGA or PGA nanoparticles decorated on the basal side showed significantly higher metabolism compared to membrane only controls for the first 14 days of culture. There was no significant difference in metabolism of the cells thereafter up to 1 month or compared with coverslip control. Significant difference *p<0.05 compared with membrane control, one-way ANOVA followed by Bonferroni post-test (n = 4). Data presented: mean (+/− standard error).

**Figure 13.**
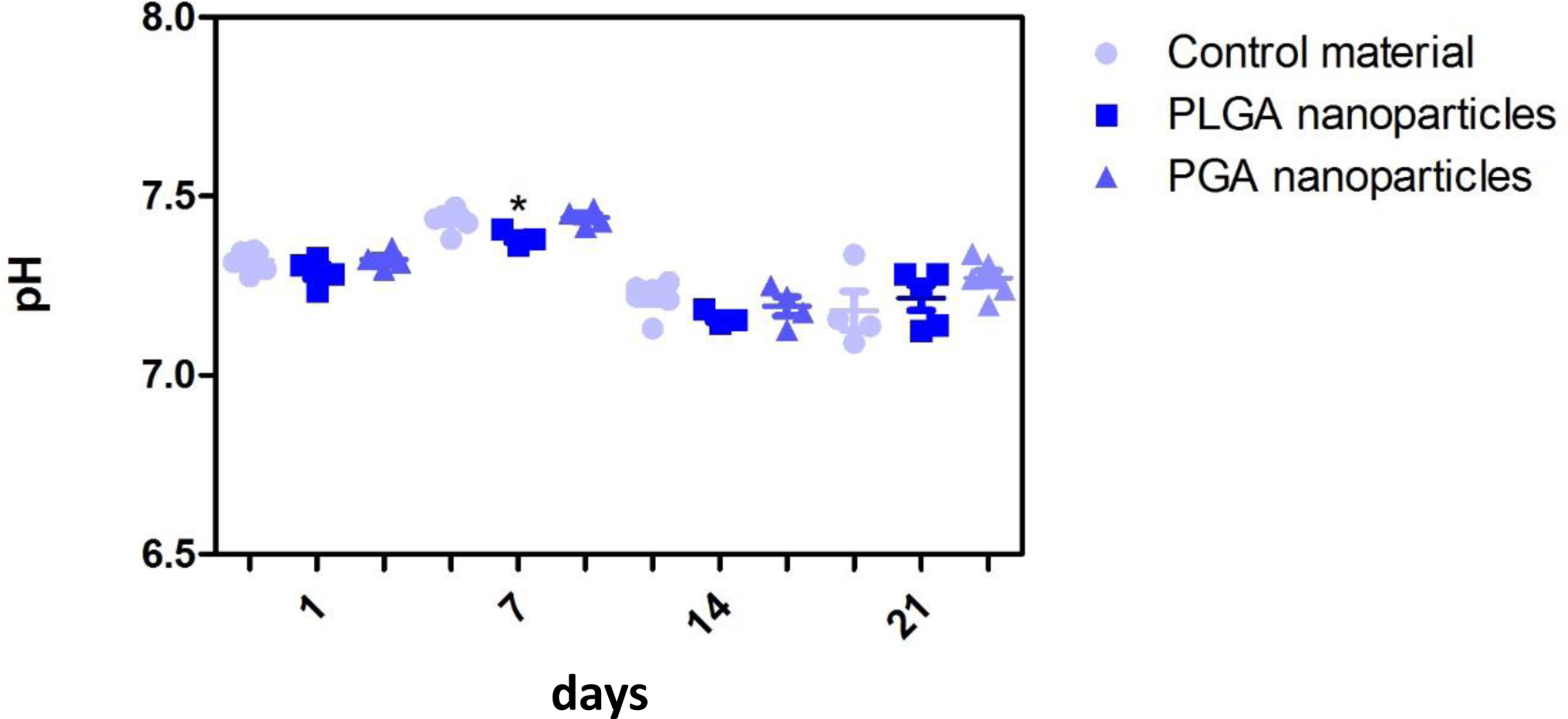
Effect of degrading nanoparticles on pH of ARPE-19 culture media exhibited little change, with PLGA exhibiting a significant decrease compared to membrane only control in week 1, which resolved thereafter. Significant difference *p<0.05 compared to PET membrane only control, one-way ANOVA followed by Bonferroni post-test (n = 3). Significant difference *p<0.05 compared to PET membrane only control, one-way ANOVA followed by Bonferroni post-test (n = 6). Data presented: mean (+/− standard error).

Although a difference in metabolism of cells and pH of their surrounding environment was observed when cultured with nanoparticles; morphology and protein expression results suggest that the cells were not adversely affected by the membrane/nanoparticle composite.

### Nanoparticle degradation and release profile

Short term degradation studies show that over 72 hours, both PLGA and PGA nanoparticles degraded releasing FITC into solution (0.1% isopropanol in dH_2_O). PLGA nanoparticles exhibited a continuous release profile up to 72 hours (figure 14A) compared to the PGA nanoparticles, which exhibited sudden burst release of the dye after 3 hours, which plateaued after 24 hours (figure 14B).

**Figure 14.**
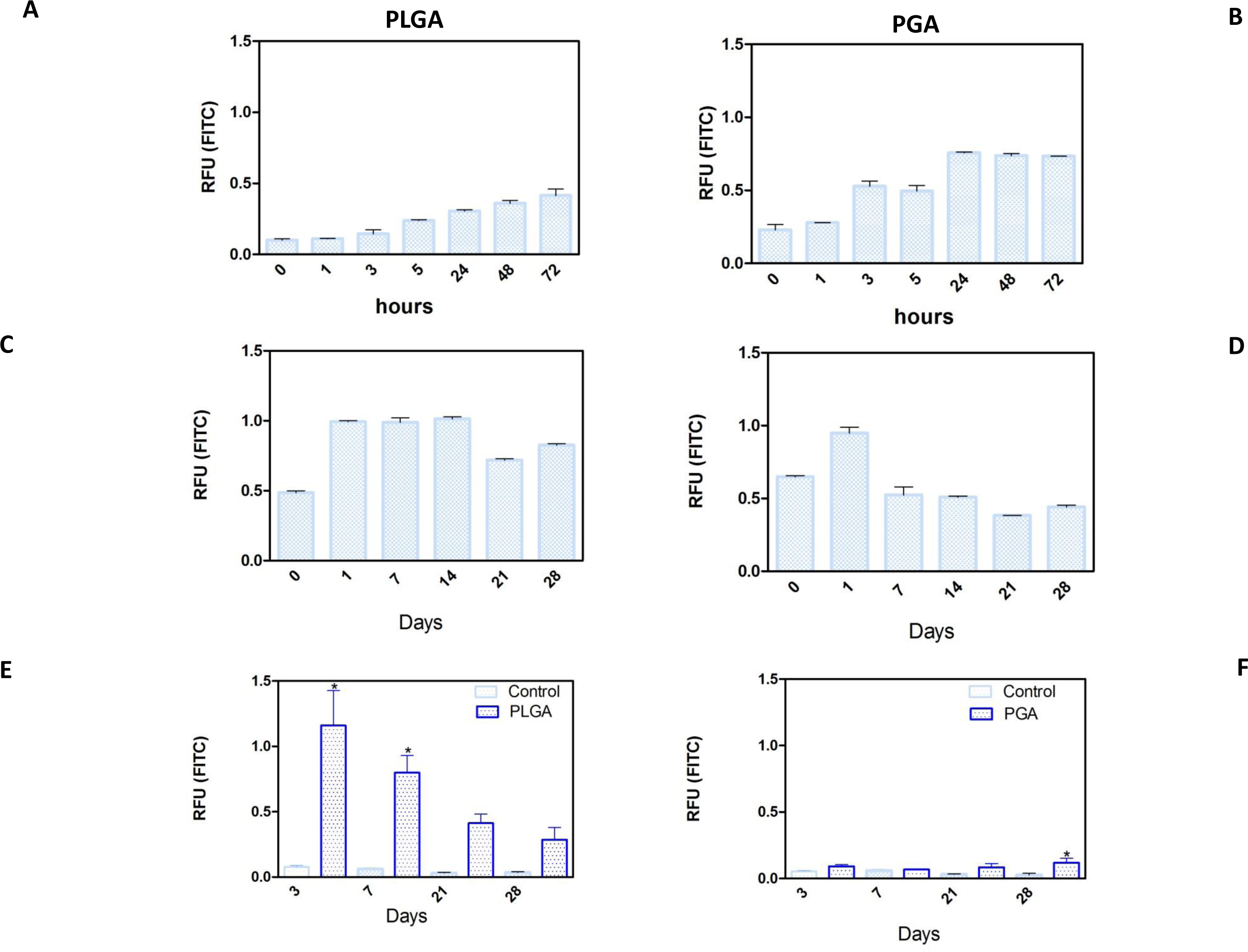
Release of FITC from PLGA and PGA nanoparticles into a solution of 0.1% isopropanol over a short period of time (**A, B**), long period of time (**C, D**) and the release of FITC into cell culture media (**E, F**) from nanoparticles electrosprayed onto the basal side of PET membrane. Significant difference *p<0.05 compared to PET membrane only control, one-way ANOVA followed by Bonferroni post-test (n = 4). Data presented: mean (+/− standard error).

Long term (up to 28 days) degradation studies showed similar results as the short-term studies with PLGA nanoparticles continuing to sustainably release FITC into solution up to 14 days, decreasing thereafter (figure 14C). PGA nanoparticles exhibited maximum release after 1 day, but exhibited considerable decrease in FITC release by day 7 onwards (figure 14D). This can be attributed to the faster degradation rate of PGA compared to PLGA and also to the higher surface area:volume ratio of significantly smaller nanoparticles of PGA, which would allow faster release of the dye, due to shorter diffusion pathway and faster degradation (Sharma, Madan et al. 2016). Observation of the resultant solution after 28 days degradation would suggest PLGA nanoparticles had released more FITC compared to PGA nanoparticles (figure S2, supplementary data), however, this could also be attributed to more FITC encapsulated. Encapsulation efficiency may explain these differences, particularly given the size difference and inter-variance in size of the produced nanoparticles; however, this was beyond the scope of this study.

Degradation studies on cell cultured PET membranes decorated with FITC encapsulated nanoparticles on the basal surface, showed PLGA nanoparticles exhibited release of FITC over 28 days. The majority of the dye release was observed up to 7 days for PLGA nanoparticles with fluorescence decreasing thereafter; however, PGA nanoparticles exhibited no significant change until after 28 days, with very little detected fluorescence in comparison to PLGA (figure 14 E-F) and upon comparison to degradation in isopropanol solution (figure 14 B). This would suggest that nanoparticle degradation in cell culture media rather than in isopropanol solution offers the nanoparticles a degree of protection from degradation. The effects of the ions, proteins, lipids and other components of the culture medium may confer a degree of stability to the nanoparticles (Moore, Rodriguez-Lorenzo et al. 2015) and is an important factor to consider in final implant design.

## Conclusion

In this article we have described the development of a composite bioactive membrane that has adequate mechanical properties, permeability and cytocompatibility. The membrane was able to maintain a monolayer of RPE cells that exhibited phenotypic microvilli structures on its apical surface while providing a sustained release of encapsulated moieties from the membrane’s basal surface.

This composite bioactive membrane exhibits the potential to act as an artificial BM and a potential treatment for atrophic AMD as a novel bioactive cell transplant substrate; to the authors’ knowledge the first of its kind to be developed as a potential treatment for atrophic AMD.

We are now focusing on the optimization and formation of co-axially electrosprayed nanoparticles to encapsulate biologically active moieties to target drusen, and have already achieved the coaxial encapsulation of FITC with Nile red as the outer shell via electrospraying (figure S3, supplementary data). Since electrospraying involves the use of compounds dissolved in solvents, care must be taken when encapsulating biologically active moieties to ensure their activity is not compromised. We have begun screening solvents for this purpose using enzyme activity agar assays (figure S4, supplementary data). Further work will then move towards using moieties such as L4-F, to target drusen.

## Author Contributions

RM performed the acquisition and undertook the analyses of the data and contributed to the writing of the manuscript. IP and SK provided critical evaluation on the progress of the work and guided the work through clinical relevance. SK also provided feedback on the manuscript. AH prepared, wrote and revised the manuscript, obtained the funding and is also leading the research.

## Abbreviations

AMD: Aged related macular degeneration
RPE: Retinal pigment epithelium
PET: poly(ethylene terephthalate)
BM: Bruch’s membrane
PLGA: poly(lactic acid-co-glycolic acid)
PGA: poly(glycolic acid)
VEGF: Vascular endothelial growth factor
FITC: Fluorescein-5-isothiocyanate
UV: Ultraviolet
DMEM: Dulbecco’s modified eagles’ media
SEM: Scanning electron microscope
DPBS: Dulbecco’s phosphate buffered saline
DAPI: 4′,6-diamidino-2-phenylindole
NBF: Neutral buffered formalin
UTS: Ultimate tensile strength
YM: Young’s modulus
WCA: Water contact angle
FTIR: Fourier-transform infrared spectroscopy
Best1: Bestrophin 1

## Acknowledgements

The authors would like to acknowledge Alison Beckett (University of Liverpool) for her help with the SEM to image the fibrous membrane and Dr Keith Arnold (Materials Innovations Factory, Liverpool) for his help with SEM in imaging the nanoparticles.

## 1 Data Availability Statement

The raw data supporting the conclusions of this manuscript will be made available by the authors, without undue reservation, to any qualified researcher.

