## supplementary data for "Optimisation of a Novel Bio-Substrate as a Treatment for Atrophic Age-Related Macular Degeneration"

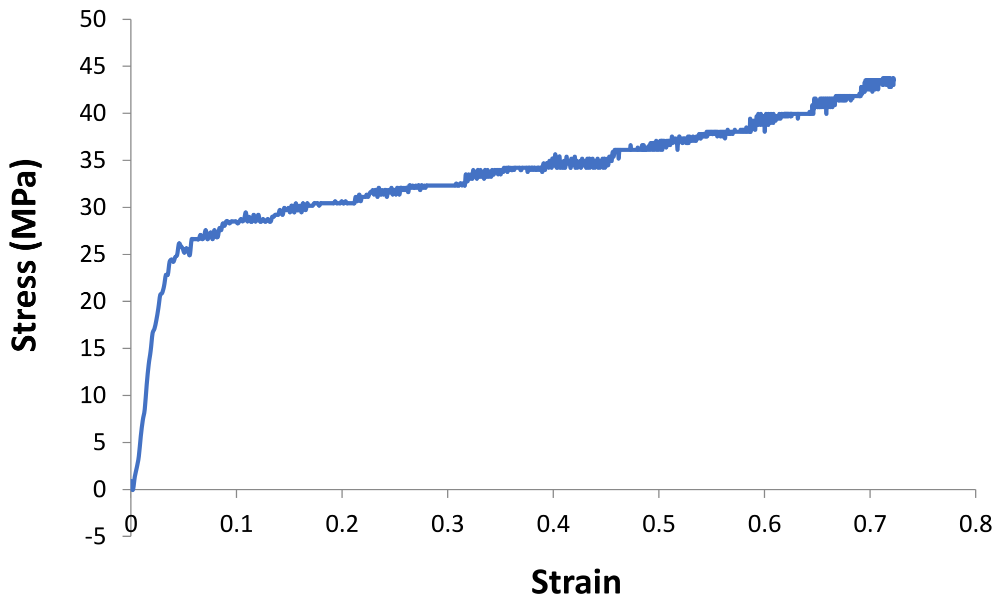


Figure S1. Representative stress vs strain profile of an electrospun membrane sample that did not achieve failure.

Table S1. Denoting the average WCA following treatment, with the media treated PET exhibiting the lowest WCA, 85.8° (+/-20). Data presented: mean (+/- standard error). (n = 6).

| **Treatment** | **Average contact angle (°)** |
|---|---|
| Control | 132.6 (+/-11.3) |
| Media | 85.8 (+/-20) |
| UV | 138.1 (+/-6.9) |
| Ethanol | 118.4 (+/-25.4) |

Table S2. Effect of degrading nanoparticles on pH of ARPE-19 culture media exhibited little change in pH, with PLGA exhibiting a significant change in week 1 and 2, which resolved thereafter. Data presented: mean (+/- standard error). (n = 6).

| **Treatment** | **pH 1 day** | **pH 1 week** | **pH 2 weeks** | **pH 1 month** |
|---|---|---|---|---|
| Control | 7.32 (+/-0.01) | 7.43 (+/-0.01) | 7.24 (+/-0.001) | 7.19 (+/-0.06) |
| 2% PLGA nanoparticles | 7.29 (+/-0.02) | 7.39* (+/-0.01) | 7.16* (+/-0.01) | 7.22 (+/-0.03) |
| 1% PGA nanoparticles | 7.32 (+/-0.01) | 7.441 (+/-0.01) | 7.193 (+/-0.03) | 7.23 (+/-0.02) |

PLGA PGA

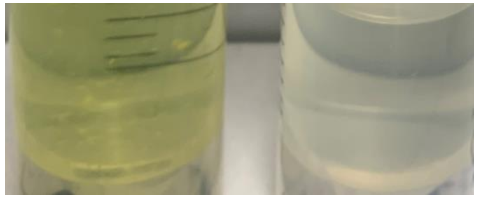

Figure S2. PLGA and PGA nanoparticles encapsulated with FITC and allowed to degrade in 0.1% isopropanol/d$H_2O$ over 28 days. Photographs suggest PLGA released more dye (see figure 14 a-b for comparison).

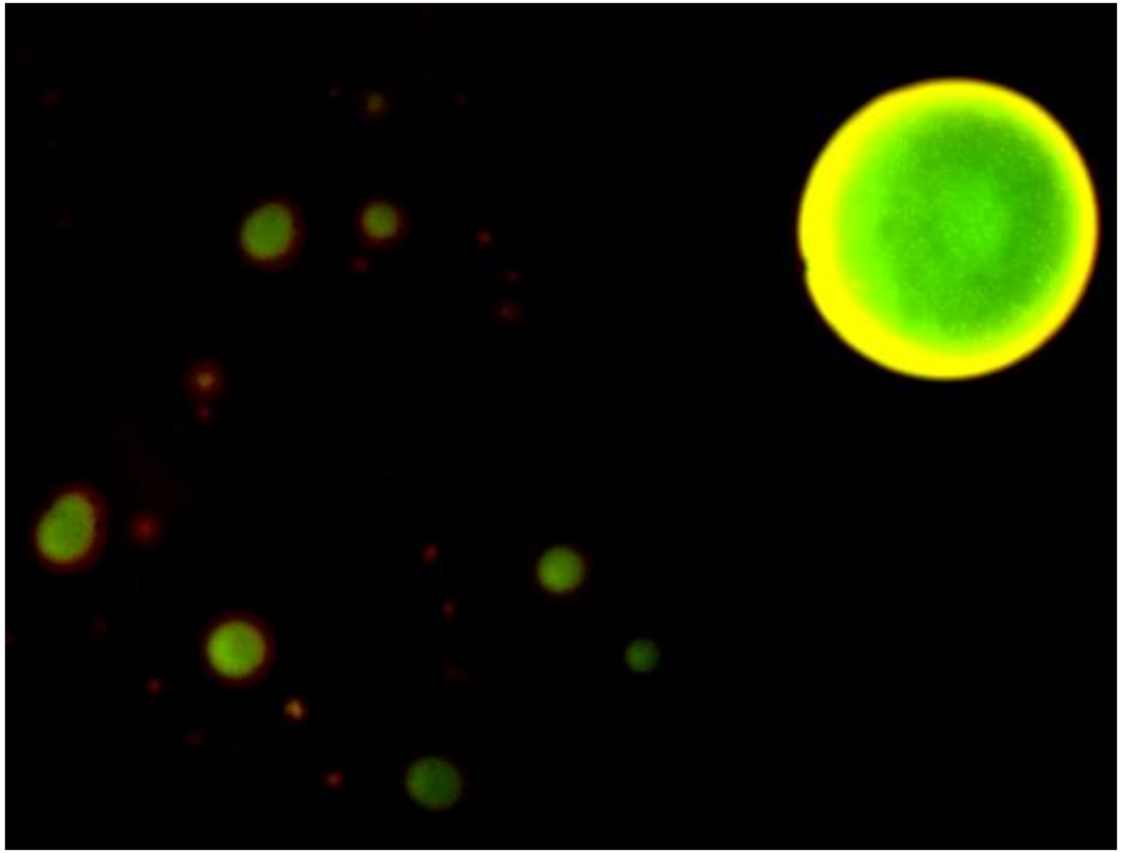

Figure S3. Confocal micrograph of coaxial electrosprayed nanoparticles with an outer shell consisting of 2% PLGA/0.01% Nile red in ethyl acetate encapsulating 0.2% PLGA/1mM FITC in acetone.

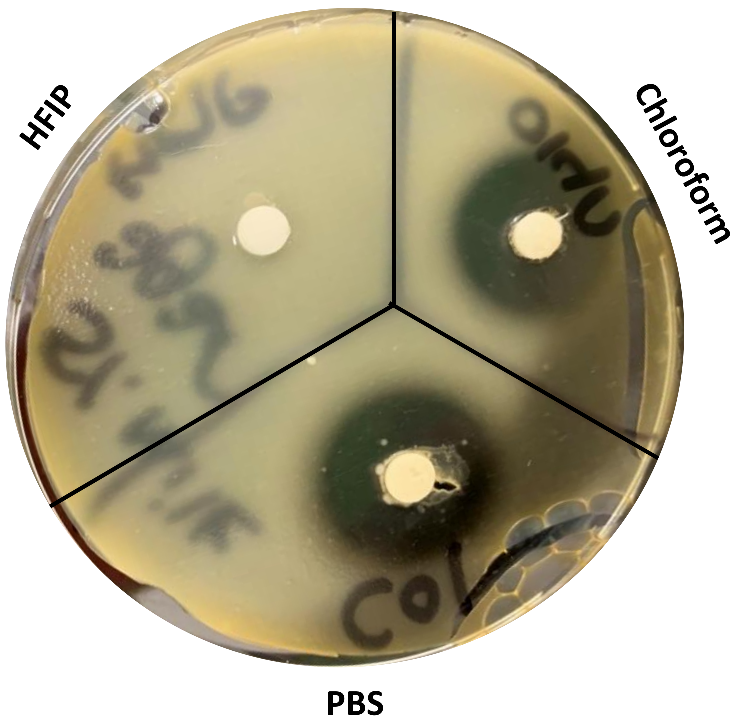


Figure S4. Enzyme activity of collagenase carried out on milk agar assay showed that HFIP denatured the enzyme whereas chloroform did not affect enzyme activity, which was comparable to collagenase dissolved in PBS.
